# Spatial goals trap hippocampal replay

**DOI:** 10.64898/2026.09.02.748857

**Authors:** Caitlin S. Mallory, John Widloski, David J. Foster

## Abstract

Hippocampal replay is proposed to support memory and planning by preferentially reactivating behaviorally significant locations^1,2^. Although learning can alter which locations are reactivated and how frequently ^3–7^, whether it also alters the dynamics by which replay propagates through the cognitive map remains unknown. To address this, we recorded large-scale CA1 activity while rats collected identical rewards at multiple spatial locations, only one of which was consistently rewarded and served as the Goal. Replay events occurred nearly twice as often during reward consumption at the Goal compared to Non-Goal sites and were accompanied by reduced inhibition. Goal-associated replays were also slower, shorter, and more locally confined. Replays originating elsewhere in the arena slowed and preferentially terminated when nearing the Goal, indicating that the Goal locally resists replay propagation regardless of where the replay was initiated. Increasing evidence suggests that hippocampal sequences arise from adaptation-like mechanisms that propagate neural activity away from recently active states^8–10^. We hypothesized that the slowing and spatial confinement of Goal-associated replay arises from reduced effective adaptation near the Goal. A recurrent network model showed that reducing adaptation strength reproduces these effects and correctly predicted increased population firing rates and weaker avoidance of recently traveled or reactivated paths. Finally, we found evidence for reduced interneuron-mediated feedback inhibition near the Goal, providing a potential circuit mechanism for locally modulating adaptation. Our findings show that learning locally reshapes the dynamics governing internally generated activity, causing replay to linger near behaviorally significant locations and providing a new mechanism for their preferential reactivation.

## Main

Hippocampal replay has been proposed to support memory-guided behavior by preferentially reactivating paths to or from behaviorally significant locations^1,2^. A classic example is the overrepresentation of reward locations: replay occurs more frequently at reward sites and replay events initiated elsewhere are biased toward remembered goals^3–6,11,12^. Thus, previous work has primarily focused on how experience shapes which locations are represented during replay. Much less is known about whether learning also changes the dynamics by which replay propagates through those representations. Here, we tested whether the learned significance of a location locally alters replay propagation through the cognitive map.

### Monitoring neural activity during reward-associated immobility

Rats (*N*=4) performed a spatial memory task within an open arena containing nine reward wells embedded in the floor (**Fig. 1A**). On odd trials, reward (0.3 mL chocolate milk) was available at a designated *Goal* well, whose location remained fixed across the session. On even trials, the same reward volume was delivered at one of the eight remaining *Non-Goal* wells, selected at random. Hereafter, immobile periods of reward consumption at the Goal and Non-Goal wells are referred to as Goal stops and Non-Goal stops, respectively. Transparent ‘jail-bar’ barriers were placed throughout the arena, which rats were required to circumvent during navigation^10^. Rats performed up to three sessions per day, separated by ∼2 hours (*N*=44 sessions total). The Goal well and barrier locations changed between sessions. Three rats performed a *delayed-reward* variant of this task in which (i) a 5–15 s delay was imposed between the end of reward consumption at one well and the filling of the next, and (ii) on Non-Goal trials, a light cue appeared adjacent to the filled well^10^.

**Fig 1.**
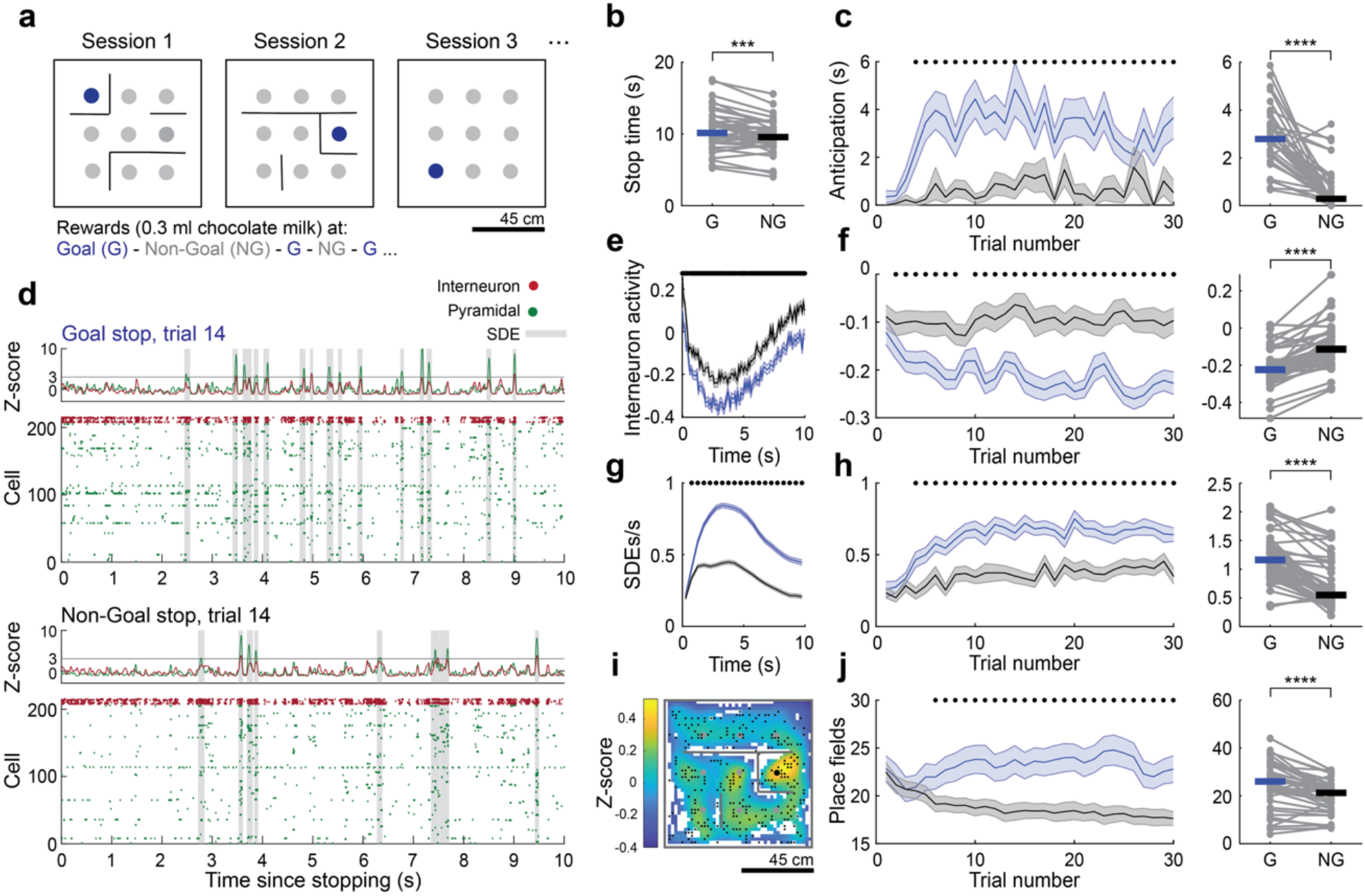
Reduced inhibition and increased population reactivation at a remembered Goal. **a**. Goal-directed navigation task setup. **b**. Average duration of Goal and Non-Goal stops (*Z*=3.6, *p*=3.7e-4, Wilcoxon signed-rank test). *N*=44 sessions from 4 rats. **c**. Anticipatory behavior near the Goal well on Goal versus Non-Goal trials as a function of trial number (left; dots indicate bins in which *p*<0.05, Wilcoxon rank-sum tests) or averaged across trials (right; *Z*=5.1, *p*=2.8e-7, signed-rank test). *N*=39 sessions from 3 rats performing the delayed reward task variation. **d**. Example raster plots showing spiking across the first 10 s of a Goal stop (top) or Non-Goal stop (bottom). Top traces show the total pyramidal and interneuron spike density. Shaded regions indicate spike density events (SDE). **e**. Interneuron spike density (z-score) during the first 10 s of Goal or Non-Goal stops. *N*=1960 Goal stops and 1939 Non-Goal stops from 44 sessions from 4 rats. Dots indicate time bins in which *p*<0.05, rank-sum tests. **f**. Interneuron spike density during Goal or Non-Goal stops as a function of trial number (left; dots indicate bins in which *p*<0.05, rank-sum tests) or averaged across trials (right; *Z*=5.6, *p*=2.8e-8, signed-rank test). *N*=44 sessions from 4 rats. **g**. Spike density event (SDE) rate across the first 10 s Goal or Non-Goal stops. *N*=1960 Goal stops and 1939 Non-Goal stops from 44 sessions from 4 rats. Dots indicate time bins in which *p*<0.05, rank-sum tests. **h**. Spike density event rate during Goal and Non-Goal stops as a function of trial number (left; dots indicate bins in which *p*<0.05, rank-sum tests) or averaged across trials (right; *Z*=5.4, *p*=5.7e-8, signed-rank test). *N*=44 sessions from 4 rats. **i**. Firing rate maps for an example session, z-scored per cell and then averaged across cells. Small black dots: center of mass of all place fields detected; Gray dots: Non-Goal wells; Large black dot: Goal well. **j**. Average number of place fields detected within 12 cm of the Goal or a given Non-Goal, as a function of trial number (left; dots indicate bins in which *p*<0.05, rank-sum tests), or averaged across trials (right, *Z*=4.6, *p*=5.2e-6, signed-rank test). *N*=44 sessions from 4 rats. \*\*\**p*<0.001, \*\*\*\**p*<0.0001. Error bars are SEM.

This task structure allowed us to compare hippocampal replay dynamics and the underlying network activity at locations associated with similar overt behavior (consumption of equivalent reward volumes), but varying in behavioral significance (only the Goal consistently predicted future reward availability). On average, reward consumption lasted ∼10 seconds during both Goal and Non-Goal stops, with slightly longer durations at the Goal (**Fig. 1B**). In the three rats performing the delayed-reward task, learning of the Goal location was evident in the development of anticipatory behavior: over the first few trials, rats increasingly lingered near the Goal before reward delivery, and this behavior was stronger on Goal versus Non-Goal trials (**Fig. 1C**).

During task performance, we recorded spiking activity from large ensembles of dorsal CA1 cells, classified as pyramidal cells (mean ± standard error of the mean (SEM) = 211 ± 9 cells per session) and interneurons (12 ± 1 cells per session) based on waveform and firing rate (**Fig. 1D**, Methods). We identified population spike density events (SDEs) as transient increases in pyramidal population activity during immobility (Methods). Within these events, hippocampal activity represented individual locations or sequences of locations, which we refer to collectively as replay.

### Learning alters hippocampal population dynamics at the Goal

We first asked whether learning altered the local hippocampal network state in which replay occurred. Local inhibition in the hippocampus strongly regulates the temporal dynamics of pyramidal spiking and the expression of population reactivation events^6,13–15^, and reduced inhibition at reward sites has recently been shown to promote reactivation events^6^. We therefore asked whether interneuron firing rates differed throughout Goal and Non-Goal stops. We computed interneuron activity outside SDEs to isolate background inhibition from interneuron participation in reactivation events (similar results were obtained when including activity within SDEs; **Fig. S1**). At both stop types, interneuron activity decreased rapidly after rats stopped to consume reward, reached a minimum after ∼3 s, and then increased again despite continued immobility and reward consumption. Despite these similar temporal profiles, interneuron firing rates were persistently lower throughout Goal stops (**Fig. 1E**).

We next examined how quickly this reduction in interneuron activity emerged within a session. A significant reduction in interneuron activity at the Goal well was detectable by the second trial, and interneuron activity continued to decline across subsequent trials (**Fig. 1F**). Across sessions, interneuron activity during Goal stops was consistently lower than during Non-Goal stops (**Fig. 1F**). Together, these data indicate that interneuron activity becomes selectively reduced at the Goal as animals learn its behavioral significance.

We next asked whether this reduction in inhibition was accompanied by changes in pyramidal population events. At both stop types, SDE rates rose rapidly following reward onset, peaked ∼3 s into the stopping period, and then declined (**Fig. 1G**). This temporal profile was approximately inverse to that of interneuron activity, and across stopping periods, mean interneuron activity was negatively associated with SDE rate (*r*(3897)=-0.23, *p*=2.4e-23, Pearson’s correlation, **Fig. S1**).

Strikingly, SDE rates during Goal stops were approximately twice those during Non-Goal stops, paralleling the reduction in interneuron activity. This difference was maintained throughout stopping periods and emerged early in the session (**Fig. 1 G-H**), mirroring the emergence of reduced interneuron activity. Similar results were obtained for sharp-wave ripples (**Fig. S2**). These findings extend recent work linking interneuron activity to population event rates^6^ by showing that their relationship spans multiple timescales: the two varied inversely within individual stopping periods (seconds), and their activity patterns rapidly diverged between Goal and Non-Goal sites as animals learned which location predicts future reward (minutes).

Finally, we asked whether enhanced pyramidal activity at the Goal was specific to immobility-associated population events or was also evident during movement (**Fig. 1I, Fig. S3**). Consistent with prior work^5,16^, overall population firing rates were not elevated near the Goal during movement (**Fig. S3**). However, place field density was enhanced around Goal relative to Non-Goal wells, consistent with prior reports of reward-site overrepresentation^17–23^. This spatial bias also developed over the course of the session: place field density near the Goal increased across trials, while density near Non-Goal wells gradually decreased. (**Fig. 1J**). Thus, learning increased both place field density near the Goal during movement and population event rates at the Goal during immobility.

### Replay trajectories are shorter, slower, and more locally confined during Goal stops

We next asked whether learning altered not only the frequency of replay at the Goal, but also the dynamics of activity within individual replay events. (**Fig. 2A-B**). Consistent with previous reports examining sharp-wave ripples in a similar task^24^, SDEs during Goal stops were shorter in duration (**Fig. S2**). They were also larger in amplitude and recruited more cells simultaneously compared to SDEs during Non-Goal stops (**Fig. S2**). We next used a Bayesian decoder to estimate the locations represented within each SDE. Although replays depicting extended spatial paths were observed during both Goal and Non-Goal stops, many more replays during Goal stops appeared tightly anchored to the animal’s current location (e.g., the four leftmost examples of **Fig. 2A**). Consistent with this observation, replays during Goal stops began and ended closer to the animal, spanned shorter total distances and net displacements, and propagated more slowly through space (**Fig. 2B**).

**Fig 2.**
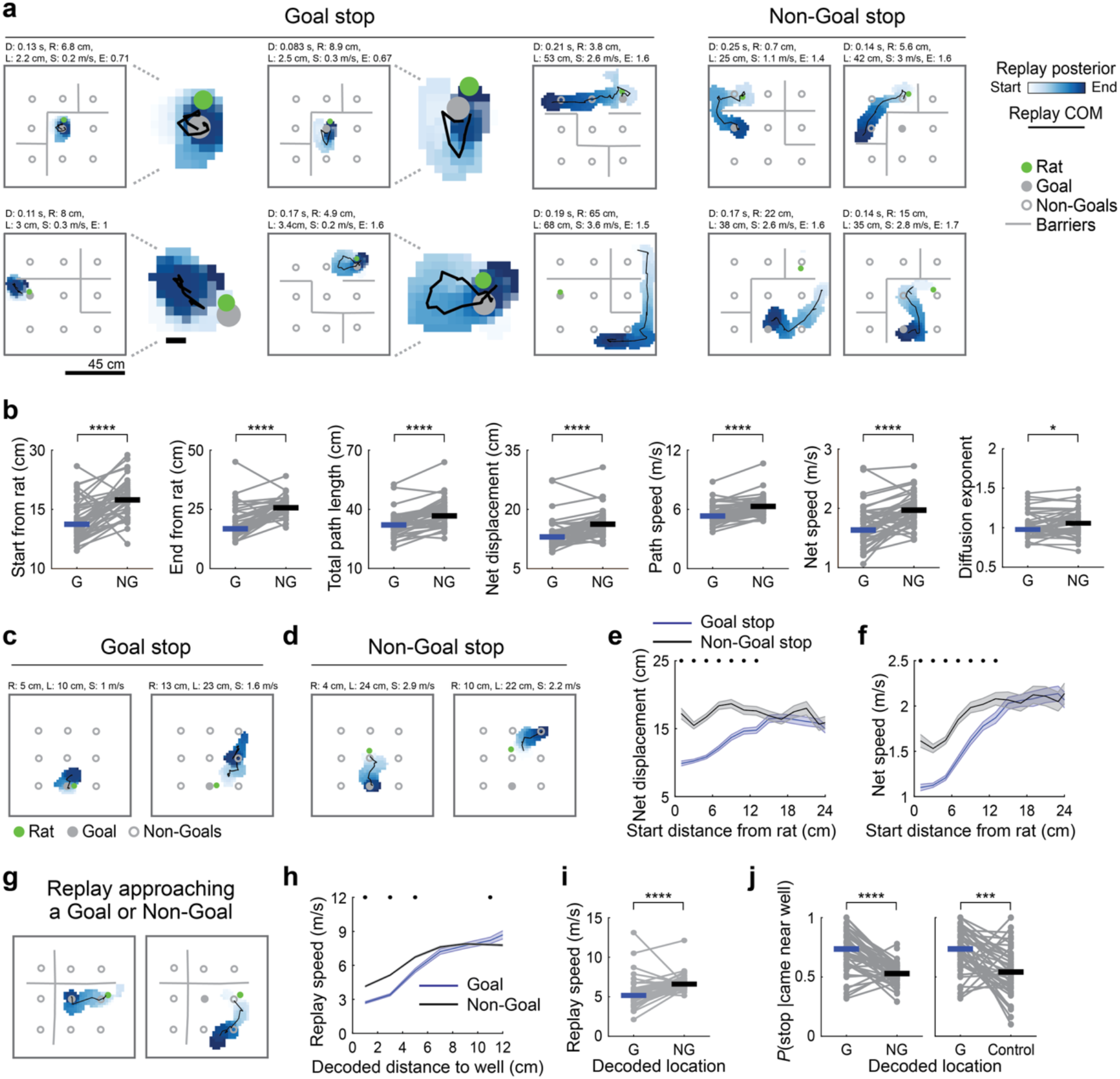
Replay trajectories slow and terminate near Goal locations. **a**. Example replays during Goal stops (left) or Non-Goal stops (right). Many replays during Goal stops represent locations close to the animal. The four examples at left include zoomed-in views. Scale bars: 45 cm. **b**. Replay properties differed during Goal and Non-Goal stops. Replays during Goal stops started closer to the rat (start from rat; *Z*=-4.4, *p*=1.3e-5), ended closer to the rat (end from rat; *Z*=-4.8, *p*=1.4e-6), spanned a shorter integrated distance (total path length; *Z*=-4.7, *p*=2.7e-6), spanned a shorter Euclidean distance (net displacement; *Z* =-5.1, *p*=4.1e-7); progressed more slowly (path speed; *Z*=-4.6, *p*=3.8e-6), progressed away from the start location more slowly (net speed; *Z*=-4.7, *p*=3.2e-6), and were less superdiffusive (diffusion exponent; *Z*=-2.3, *p*=0.024). Signed-rank tests. Plots show session averages from 44 sessions from 4 rats. **c**. Examples of replays during a Goal stop that started close to (left) or farther from the rat (right). Abbreviations; R: replay start distance from the rat; L: replay net displacement; S: Replay net speed. **d**. As in [c], but for replays occurring during a Non-Goal stop. **e**. Replay net distance as a function of the replay’s starting distance from the rat. Blue and gray traces show the mean ± SEM across all replays at Goal or Non-Goal stops, respectively. Dots indicate distance bins for which *p*<0.05 (rank-sum tests). **f**. Replay net speed as a function of the replay’s starting distance from the rat. Blue and gray traces show the mean ± SEM across all replays at Goal or Non-Goal stops, respectively. Dots indicate distance bins for which *p*<0.05 (rank-sum tests). **g**. Example replays during a Non-Goal stop that pass near the Goal (left) or another Non-Goal (right). **h**. Instantaneous replay speed as a function of the decoded distance to the nearest well (blue: Goal, gray: Non-Goal). Local representations within 12 cm of the animal were removed from analysis. Plots show mean ± SEM across decoding frames. Dots indicate distance bins for which *p*<0.05 (rank-sum tests). **i**. Instantaneous speed of remote replay within 12 cm of the Goal or Non-Goal. Remote replay near the Goal was slower than remote replay near a Non-Goal (*Z*=-4.5, *p*=7.1e-6, signed-rank test). *N*=43 sessions containing nonlocal replay in each group. **j**. Conditional probability of stopping near a well given that a remote replay entered within 12 cm of it. Left: replays were more likely to stop near the Goal versus a Non-Goal well (*Z*=4.7, *p*=3.1e-6, signed-rank test). *N*=43 sessions containing at least one remote replay entering each well type. Right: replays were more likely to stop near the Goal versus a distance-matched Non-Goal control well located at a similar distance from the rat as the Goal (*Z*=3.7, *p*=0.000024, signed-rank test). *N*=43 sessions containing at least one remote replay entering each well type. \**p*<0.05, \*\**p*<0.01, \*\*\**p*<0.001, \*\*\*\**p*<0.0001.

Replay trajectory dynamics can be described along a continuum from subdiffusion (constrained trajectories) to Brownian diffusion (random walk) to superdiffusion (directed motion)^9,25^. We therefore asked whether Goal-associated replay exhibited more constrained dynamics. To quantify replay diffusivity, for each session we separately concatenated all replays at the Goal or Non-Goal wells and measured the relationship between the temporal separation of all pairs of decoding windows and the distance between their decoded locations. We fit a power-law exponent to this relationship, with exponents near 1 indicating Brownian diffusion, values below 1 indicating subdiffusion, and values above 1 indicating superdiffusion. The mean exponent was significantly lower during Goal stops, with Goal-associated replays more strongly resembling Brownian motion and Non-Goal replays exhibiting modest superdiffusion (mean ± SEM; Goal: 1.02 ± 0.024, Non-Goal: 1.07 ± 0.026; **Fig. 2B**). Although exponent values varied with replay inclusion criteria, Goal stops consistently exhibited lower exponents than Non-Goal stops (**Fig. S4**). The diffusion coefficient, reflecting the rate of spatial spread, was also reduced during Goal stops (**Fig. S4**). Thus, learning was associated not only with more frequent replay at the Goal, but with altered propagation of individual replay trajectories.

### Replay trajectories slow and terminate at the Goal

We hypothesized that these differences in replay dynamics reflect the learned significance of the spatial locations represented during replay, rather than (or in addition to) the animal’s current behavioral state (i.e., preparing for exploration versus Goal-directed movement). If the Goal locally alters replay propagation, we reasoned that differences between Goal and Non-Goal stops should be largest when replay represents locations near the animal’s current reward well and diminish as replay moves farther away. To test this prediction, we examined replays beginning at different distances from the rat during Goal and Non-Goal stops (**Fig. 2C–D**), where the beginning of the replay was defined as the first decoded location within the SDE. Replays beginning near the animal (<12 cm; *proximal*) during Goal stops spanned shorter net distances than proximal replays during Non-Goal stops (**Fig. 2E**). In contrast, replays beginning farther from the rat (12**–**24 cm; *distal*) spanned similar distances regardless of stop type. Replay speed showed a similar spatial dependence: although proximally initiated replays were slower at both stops, they were especially slow during Goal stops (**Fig. 2F**). Thus, differences in replay dynamics between Goal and Non-Goal stops were greatest near the animal’s current location, consistent with a localized effect of the Goal on replay propagation.

To test this idea independently of the animal’s current location, we next examined remote replay (representations >12 cm from the animal) during Non-Goal stops. We measured the instantaneous speed of remote replay trajectories as they approached either the Goal or Non-Goal wells (**Fig. 2G**). Replay slowed significantly more near the Goal than near Non-Goal wells (**Fig. 2H-I**), an effect that contributed to overrepresentation of the Goal during replay (**Fig. S2**). We next asked whether this slowing was accompanied by an increased probability of replay termination.

Restricting analyses to Non-Goal stops, we computed the conditional probability that a replay terminated near a well given that it passed within 12 cm of it. Replays were significantly more likely to terminate near the Goal than near Non-Goal wells, including a control well equidistant from the rat’s current location as the Goal (**Fig. 2J**). These results demonstrate that the Goal exerts a localized influence on replay dynamics, slowing propagation and increasing the likelihood that trajectories terminate nearby. Moreover, these changes persist when the Goal is represented remotely, suggesting that they reflect the local reorganization of hippocampal dynamics rather than differences in the animal’s behavioral state.

### A recurrent network model with reduced adaptation strength reproduces Goal-associated replay dynamics

We further asked what mechanism could account for the altered propagation of replay near the Goal. Recent work suggests that hippocampal replay can emerge in recurrent networks endowed with spike-frequency adaptation^8–10^. In these models, which rely on continuous attractor dynamics to support a stable activity bump in the network, adaptation biases movement of the bump away from recently active states, generating sequential activity. Stronger adaptation tends to produce faster, more directed trajectories, whereas weaker adaptation produces slower, more spatially constrained trajectories^9^. Given that replays near the Goal were slow and spatially constrained, we hypothesized that effective adaptation strength may be locally reduced near the Goal.

We simulated hippocampal place cell activity using a continuous attractor network in which cells exhibited spike-frequency adaptation characterized by a recovery time constant (τ_a_) and adaptation strength (w_a_) that was spatially uniform within each simulation but varied across simulations (**Fig. 3A**). For simplicity, we modeled exploration on a 1D circular track, which included a series of movement bouts and stopping periods. Sequences were generated throughout the simulation by pulsing spatial inputs related to the animal’s location into the network and then allowing recurrent dynamics to take over (Methods). Longer inter-pulse intervals were used to simulate replay during stopping periods, and shorter inter-pulse intervals were used to simulate neuronal sequences during movement (theta sequences^26,27^). We found that reducing adaptation strength while keeping the time constant fixed caused replay sequences to propagate more slowly, resulting in shorter spatial trajectories (**Fig. 3B-C**). Movement-associated theta sequences were also shorter under reduced adaptation, reflecting slower propagation over the fixed sequence duration (**Fig. S5**). Consistent with this prediction, Goal-associated theta sequences were both shorter and slower in the empirical data (**Fig. S5**).

**Fig 3.**
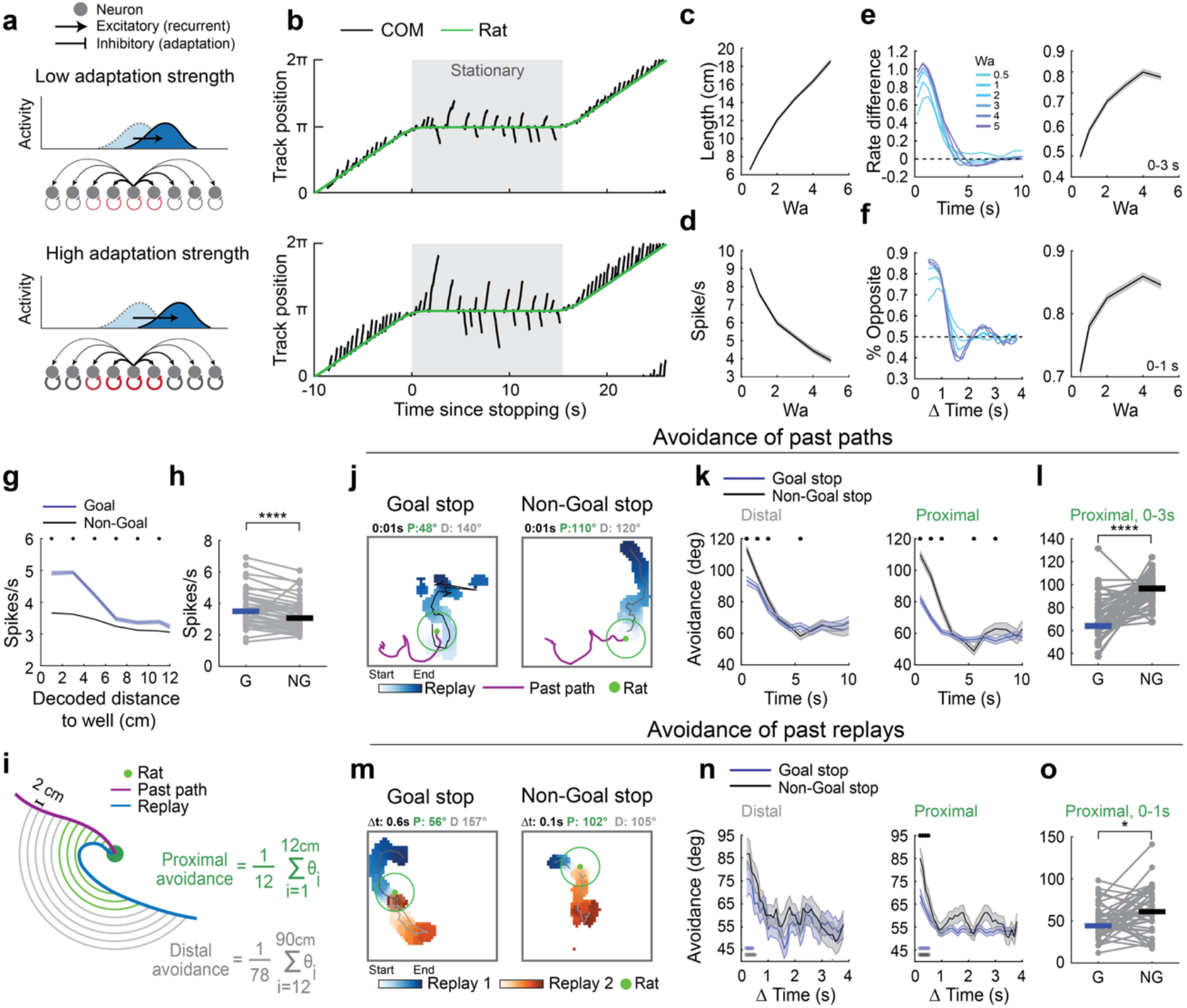
A network model with reduced adaptation strength reproduces Goal-associated replay dynamics. **a**. Recurrent network model. Cells received recurrent excitatory input from other pyramidal cells and an adaptive inhibitory input that depended on each cell’s recent spiking history. Top and bottom panels illustrate network dynamics with varying adaptation strengths (*W_a_*). For the adaptive input, stroke color illustrates the temporal decay of inhibition (red: active inhibition), and stroke weight illustrates the strength of the inhibition (*W_a_*). **b**. Population activity during simulated track traversal with lower or higher adaptation strength (top: *W_a_* = 1, bottom: *W_a_* = 4). Sequences occurring when the rat was stationary (gray) were considered replay events and quantified further. The circular track is linearized for visualization. **c**. Replay length (net displacement from start to end location) increased with adaptation strength (*r*(4)=0.997, *p*=1.7e-5; Pearson’s correlation). Plot shows mean ± SEM replay length from 100 model runs each with *W_a_* = 0.5, 1, 2, 3, 4, or 5. **d**. Mean firing rate during replay events decreased with increasing adaptation strength (*r*(4)=0.98, *p*=1e-3; Pearson’s correlation). **d**. Replay–past avoidance increased with adaptation strength. Left: replay–past avoidance as a function of time since stopping for models with different adaptation strengths. Replay–past avoidance was quantified as the difference in the rate of replay trajectories ahead of the animal versus behind the animal, such that larger values indicate stronger avoidance of recently traversed paths. Traces show mean replay–past avoidance for models with *W_a_*=0.5, 1, 2, 3, 4, or 5. Right: replay–past avoidance in the first 3 s after stopping increased with adaptation strength (*r*(4)=0.94, *p*=5e-3, Pearson’s correlation). Plot shows mean ± SEM from 800 model runs each. **f**. Replay–replay avoidance increased with adaptation strength. Left: replay–replay avoidance plotted as a function of the time between replay events for models with different adaptation strengths. Replay–replay avoidance was quantified as the percentage of replay pairs expressing opposite trajectory content (one replay in front of the animal and one replay behind the animal), such that larger values indicate stronger avoidance of recently replayed trajectories. Traces show mean replay–replay avoidance for model runs with *W_a_*=0.5, 1, 2, 3, 4, or 5. Right: replay– replay avoidance for replay pairs occurring within 1 s of one another increased with adaptation strength (*r*(4)=0.88, *p*=0.02, Pearson’s correlation). **g**. Empirical data showing the average firing rate of neurons as remote replays approached either the Goal or a Non-Goal well (mean ± SEM across decoding frames). Note elevated firing rates near the Goal, as predicted by models with lower adaptation strength. **h**. Firing rates during remote replay within 12 cm of the Goal were elevated relative to firing rates during remote replay within 12 cm of a Non-Goal (*Z*=3.4, *p*=0.00066, signed-rank test). *N*=43 sessions containing nonlocal replay near each well type. Local representations within 12 cm of the rat’s current location were excluded from analysis. **i**. Schematic illustrating calculation of replay–past avoidance and replay–replay avoidance in the empirical data. Avoidance was quantified as the mean absolute angular displacement between a replay trajectory and the past path (or past replay trajectory) at increasing distances from the animal. To calculate angular displacement, circles of increasing radius were centered on the animal, and the angle between the two trajectories was measured from the arc separating their intersections with each circle. Larger angular displacements indicate stronger avoidance. Avoidance was calculated separately for trajectory segments near the animal (proximal, green shading) and far from the animal (distal, gray shading). **j**. Example replays occurring during Goal or Non-Goal stops, with the animal’s past path shown in purple. Segments within the green circle are considered proximal and those outside considered distal. The time since stopping, proximal replay–past avoidance (*P*), and distal replay–past avoidance (*D*) are indicated above each example. **k**. Proximal (right) and distal (left) replay–past avoidance as a function of time since stopping (left). Traces show the mean ± SEM across all replays occurring during Goal (blue) or Non-Goal (gray) stops. Dots indicate time bins for which *p*<0.05 (rank-sum tests). **l**. Session-averaged proximal replay–past avoidance for replays occurring within the first 3 s after stopping. Proximal past avoidance was reduced on Goal versus Non-Goal stops (*Z*=-4.9, *p*=8.4e-7, signed-rank test). *N*=44 sessions. **m**. Example replay pairs during a Goal or Non-Goal stop. Segments within the green circle were classified as proximal and those outside as distal. The time between replays (Δt), proximal replay–replay avoidance (*P*), and distal replay–replay avoidance (*D*) are indicated above each example. **n**. Proximal (right) and distal (left) replay–replay avoidance as a function of time between replay events. Blue and gray traces indicate the mean ± SEM for replay pairs occurring during Goal or Non-Goal stops, respectively. Dots at top indicate time bins for which Goal and Non-Goal replay–replay avoidance differed (*p*<0.05, rank-sum tests). Dots at bottom indicate time bins in which replay–replay avoidance at the Goal (blue) or Non-Goal wells (grey) exceeded the 97.5th percentile of a null distribution generated by permuting time differences between replay pairs. **o**. Session-averaged proximal replay–replay avoidance for replay pairs separated by less than 1 s. Proximal replay–replay avoidance was reduced on Goal versus Non-Goal stops (*Z*=-2.3, *p*=0.019, signed-rank test). *N*=44 sessions. \**p*<0.05, \*\*\*\**p*<0.0001

Reducing adaptation also produced additional experimentally testable predictions. If Goal-associated replay dynamics indeed reflect reduced effective adaptation, these predictions should also be evident in the empirical data. First, weaker adaptation increased the local persistence of activity, resulting in higher instantaneous population firing rates during replay (**Fig. 3D**). Consistent with this prediction, population firing rates during remote replay (>12 cm from the rat) increased as trajectories approached the Goal and were higher near the Goal than near Non-Goal wells (**Fig. 3G-H**). The model also predicted that weaker adaptation would reduce the influence of recent activity on replay content. Adaptation suppresses recently active representations, causing replay trajectories to transiently avoid the animal’s most recent path for ∼3 s after stopping (replay– past avoidance) and reducing the likelihood that two replay events occurring within ∼1 s of one another represent the same trajectory (replay–replay avoidance)^8^. We previously showed that the temporal dynamics of these avoidance effects are largely governed by the adaptation time constant; here, we asked whether adaptation strength controls their magnitude. We quantified replay–past avoidance as the difference in the rates of replay trajectories directed ahead of versus behind the animal’s recent path. Likewise, we quantified replay–replay avoidance as the percentage of replay pairs expressing opposite paths. In the model, reducing adaptation strength reduced the magnitude of both replay–past and replay–replay avoidance without altering their timescales (**Fig. 3E-F**).

### Replay near the Goal shows reduced avoidance of recent activity

We next tested these avoidance predictions in the empirical data. In the 2D environment, replay– past avoidance was quantified by the angular separation between replay and the animal’s most recent path, and replay–replay avoidance was quantified by the angular separation between pairs of replay trajectories. Larger angles indicate greater avoidance (**Fig. 3I**). Because our earlier results showed a spatially localized effect of the Goal on replay propagation, we quantified avoidance separately for trajectory segments proximal (<12 cm) and distal (>12 cm) to the current reward well (**Fig. 3I**). Proximal replay–past avoidance was strongly reduced during Goal versus Non-Goal stops, whereas distal avoidance was only modestly reduced (**Fig. 3J-L**). Likewise, proximal replay–replay avoidance was reduced during Goal stops relative to Non-Goal stops (**Fig. 3M-O**). Together, these results indicate that recently active representations exert less influence over subsequent replay near the Goal, as predicted by models with reduced adaptation strength. Also consistent with the model, while the magnitudes of replay–past and replay–replay avoidance were reduced at the Goal, their temporal decay profiles were similar across Goal and Non-Goal stops.

### Effective transmission between pyramidal cells and interneurons is reduced near the Goal

Finally, we asked what biological mechanisms might underlie the apparent reduction in adaptation near the Goal. Given that we observed location-specific reductions in interneuron activity (**Fig. 1E–F**), we first examined the balance between pyramidal and interneuron activity near the Goal. The inhibition-to-excitation ratio was reduced both during Goal stops relative to Non-Goal stops (**Fig. 4A**) and during remote replay of the Goal (**Fig. 4B-C**). Goal stops were associated with both increased excitation and reduced inhibition, whereas remote representations of the Goal exhibited elevated excitation without a corresponding increase in inhibition (**Fig. S6**). These findings suggested that pyramidal cells representing the Goal recruit feedback inhibition less strongly.

**Fig. 4.**
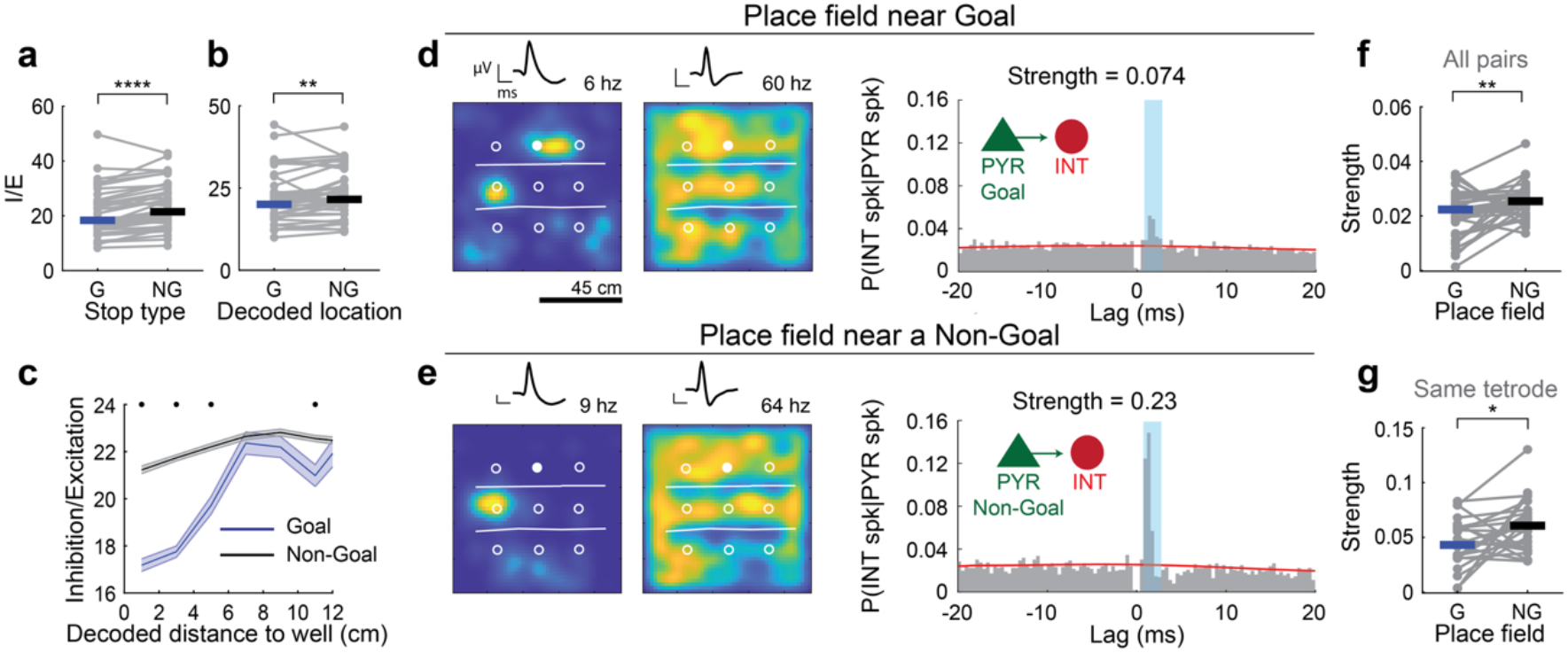
Effective transmission between pyramidal cells and interneurons is reduced near the Goal. **a**. The inhibition-to-excitation ratio (I/E) was reduced during Goal stops versus Non-Goal stops (*Z*=-4.9, *p*=8.4e-7, signed-rank test). *N*=44 sessions. **b**. The inhibition-to-excitation ratio (I/E) during remote replay within 12 cm of the Goal was reduced relative to remote replay within 12 cm of a Non-Goal well (*Z*=-2.7, *p*=0.0061, signed-rank test). *N*=43 sessions containing nonlocal representations near each well type. Local representations within 12 cm of the rat’s current location were excluded from analysis. **c**. Inhibition-to-excitation ratio (I/E) as a function of the distance between the replay and the nearest well. Traces show the mean ± SEM I/E ratio for all replay near the Goal (blue) or Non-Goal wells (gray). Local representations (<12 cm from the animal) were excluded. Dots at top indicate distance bins for which I/E differed between Goal and Non-Goal replay (*p*<0.05, rank-sum tests). **d**. An example putative monosynaptic pyramidal–interneuron (PYR–INT) pair in which the pyramidal cell fired maximally near the Goal. Heat maps show the spatial firing rate maps for the pyramidal cell (left) and interneuron (right), respectively. Scale bar: 45 cm. The corresponding spike waveforms and peak firing rates (scalebars: 0.33 ms, 50 μV) are indicated above. Open white circles: Non-Goal wells. Filled white circles: Goal well. White lines: barriers. Right: cross-correlogram normalized by the number of presynaptic PYR spikes (probability of INT spiking per PYR spike per bin). The red trace shows a slow-timescale Poisson baseline estimate. Spike transmission strength (Ptrans) was calculated as the area of the baseline-subtracted CCG within the monosynaptic window (blue; 0.8–2.8 ms). **e**. As in (d), but for an example putative monosynaptic PYR–INT pair in which the pyramidal cell fired maximally near a Non-Goal well. **f**. Spike transmission strength was reduced for PYR–INT pairs in which the pyramidal cell fired maximally near the Goal, relative to pairs in which the PYR cell fired maximally near a Non-Goal well (*Z*=-3.0, *p*=0.0026, signed-rank test). *N*=39 sessions containing at least one PYR–INT pair in each category. **g**. As in (F), but only considering PYR–INT pairs recorded on the same tetrode (*Z*=-2.4, *p*=0.015, signed-rank test). *N*=26 sessions containing at least one PYR–INT pair in each category. \**p*<0.05, \*\**p*<0.01, \*\*\*\**p*<0.0001

To test this hypothesis more directly, we identified putative monosynaptically connected pyramidal cell–interneuron (PYR–INT) pairs and quantified spike transmission probability^15,28^ (**Fig. 4D-E**; Methods). Transmission probability was lower for pairs in which the pyramidal cell’s spatial tuning was concentrated near the Goal (**Fig. 4F**). Because transmission probability is expected to be strongest for locally recorded pairs^15^, we additionally restricted analysis to PYR– INT pairs recorded on the same tetrode. These pairs exhibited higher overall transmission probabilities, but transmission remained significantly reduced for Goal-tuned pairs (**Fig. 4G**). Together, these findings implicate reduced local feedback inhibition near the Goal as a potential circuit mechanism for weakening adaptation-like dynamics and altering replay propagation.

## Discussion

Replay is increasingly understood as a dynamical phenomenon. Reactivation events span a continuum from stationary representations to rapidly propagating trajectories^29^, and their structure varies across behavioral states^24,25,30^, experience^31^, and even individual immobility periods^8,32^. Our results demonstrate that learning the significance of a location alters the local dynamics by which replay propagates through the region. As a result, replays initiated at the Goal or passing through it linger locally rather than extending across the cognitive map. This local slowing thus promotes the overrepresentation of behaviorally significant locations^5,33^, while potentially operating alongside processes that bias replay toward goal locations from a distance^5,34^.

We propose that these effects arise from modulation of adaptation-like dynamics within hippocampal circuits. In recurrent network models, spike-frequency adaptation generates sequential activity by biasing activity away from recently visited states^8,9,35–38^. Strong adaptation promotes rapid propagation of activity, whereas weaker adaptation produces more spatially constrained dynamics. Our findings that replay near the Goal was slower, more spatially constrained, and less dependent on recent activity point to reduced effective adaptation in this region. Flexible adaptation processes have previously been proposed to shape neural sequence dynamics across brain states (e.g., awake immobility versus sleep^9^), and behavioral timescales (modulation across individual theta cycles^28^). Our findings extend this framework by suggesting that adaptation-like dynamics can also vary across the represented space, allowing different regions of the same cognitive map to support distinct modes of sequence propagation.

We further asked what circuit mechanisms might contribute to the apparent reduction in adaptation-like dynamics near the Goal. Adaptation can arise from multiple biological factors, including intrinsic membrane conductances such as the afterhyperpolarization^39,40^ or short-term synaptic depression^41^. In addition, local inhibitory feedback can regulate how strongly adaptation shapes network dynamics^42,43^. Our data support a role for this latter mechanism. Transmission probability between putative pyramidal cells and interneurons was reduced when the pyramidal cell was Goal-tuned, consistent with weaker recruitment of local inhibitory feedback, and interneuron firing rates were also lower throughout Goal stops compared to Non-Goal stops^6^. Notably, the temporal profile of interneuron activity was remarkably similar across stop types, differing primarily in gain. This parallels our modeling results, in which Goal-like replay emerged by reducing adaptation strength while preserving its recovery time constant. Likewise, replay avoidance of recent activity was selectively reduced at the Goal while decaying with a similar time course. These observations raise the possibility that learning modulates the strength of local inhibitory feedback without altering its temporal dynamics.

Here, we show that learning creates spatially localized changes in hippocampal dynamics such that the rules governing internally generated activity vary across the cognitive map. As a result, replay lingers at behaviorally significant locations rather than propagating uniformly through space. More broadly, local modulation of neural dynamics may represent a general mechanism by which experience prioritizes behaviorally significant representations.

## Materials and Methods

### Experimental model and subject details

All experimental procedures were performed in accordance with the University of California Berkeley Animal Care and Use Committee and US National Institutes of Health guidelines. Subjects were male Long-Evans rats (*Rattus norvegicus*; 3-9 months old, 450-550 g; Charles River Laboratories). Rats were housed in a humidity- and temperature-controlled facility with a 12 h light-dark cycle. Before starting experiments, rats from the same breeding cohort were co-housed in pairs. Rats were single-housed at the start of experiments. All rats were implanted with microdrives containing multiple tetrodes targeting the hippocampal CA1 pyramidal layer (see Drive design and surgery). Data presented here include re-analysis of previously published data (Cohort 1: 3 rats from^10^, Cohort 2: 1 rat from^8^(Mallory et al., 2025); this animal was injected with AAV encoding Jaws bilaterally in medial entorhinal cortex, but only data from control sessions (no-laser) were analyzed here).

### Drive design and surgery

Rats were implanted with microdrives, as previously described^10^. Briefly, rats were anesthetized via 5% isoflurane inside an induction chamber and then transferred into a stereotaxic apparatus where they were maintained at 1-3% isoflurane. Rats received injections of buprenorphine (0.1 mg/kg), atropine (5mg/kg), and cefazolin (5mg/kg) at the start of surgery. A microdrive array weighing 40-50 g and containing 40-64 independently movable tetrodes was secured above the dorsal CA1 region of hippocampus (position from bregma: AP -4.1 mm, ML ± 2.7 mm) using bone screws and dental cement. Before implantation, tetrodes (platinum iridium) were gold-plated to achieve impedance of 150-300 MΩ. Tetrodes were gradually lowered into the CA1 pyramidal layer over the course of 2-6 weeks, which was identified by the presence and polarity of sharp-wave ripples. The rats were allowed a week of recovery, after which food restriction and behavioral training resumed.

### Behavioral task and apparatus

Four rats performed a navigation task within a raised, square maze (90 × 90 cm). The floor contained 9 hidden reward wells evenly spaced on a 3 × 3 grid, with an interwell distance of 23 cm along each dimension. The task consisted of alternate trials of goal-directed navigation to a fixed, unmarked Goal well and navigation to one of 8 randomly selected Non-Goal wells. For each session, the position of 6 transparent ‘jail’ barriers was selected pseudorandomly. Six sessions contained no barriers. The Goal well location and the order of Non-Goal wells changed between sessions. Rats performed between 1 and 3 sessions per day, beginning a few weeks after surgery. 3 rats performed a variation of this task in which a) a visual cue marked the baited Non-Goal wells and b) a variable 5-15 delay was imposed between the end of reward consumption at one well and the filling of the next well.

### Behavioral analysis

Rat position was tracked by an overhead camera and sampled at 20 Hz. Position data was determined from red and green LEDs mounted on the headstage. Position and velocity data were smoothed using a Butterworth filter (second order with a cutoff frequency of 0.1 Hz using MATLAB’s butter function). For each trial, reward consumption onset was defined as the first time the rat came within 5 cm of the filled reward well at a speed of 3 cm/s or less. The end of reward consumption was taken as the first time the rat exited the reward well at a speed of 5 cm/s or greater. In the delayed-reward task variation, in which a variable delay of 5–15 s was imposed before the filling of a well, rats often lingered near the Goal well before it was filled ^10^. We therefore quantified ‘anticipatory periods’ as times in which the rat was near a well (within 5 cm) and the smoothed velocity stayed within 3–6 cm/s.

### Freely Moving Neural Data Acquisition

Neural data were acquired using SpikeGadgets acquisition systems (sampled at 30 kHz). Spike events exceeding 50 µV were clustered automatically using MountainSort^44^. Automatically determined clusters were accepted if they passed visual inspection and met the following criteria: noise overlap < 0.03, isolation > 0.5, peak signal-to-noise ratio > 1.5. Local field potentials (LFP) were digitally filtered between 0.1 and 500 Hz and recorded at 1500 Hz. Isolated units with mean firing rates > 10 Hz and visually identified narrow waveforms were classified as putative interneurons. All remaining isolated units were classified as putative pyramidal cells.

### Spike density and sharp-wave ripple detection

Population pyramidal cell and interneuron spike density functions were computed separately by summing spikes from each cell type in 1-ms non-overlapping time bins. The local field potential (LFP), recorded from a tetrode in the pyramidal layer with visually identified sharp-wave ripples, was band-pass filtered between 150–250 Hz, and ripple amplitude was computed as the amplitude of the analytic signal obtained from the Hilbert transform. Pyramidal spike density and ripple amplitude were each smoothed with a Gaussian kernel (12.5 ms standard deviation). Analyses were restricted to periods when the rat’s speed was below 5 cm/s, and signals were z-scored within these immobility periods. Peaks in the z-scored pyramidal spike density or ripple amplitude traces that exceeded 3 SD above the mean were identified as spike density events or ripple events, respectively. The start and end of each event were defined as the time points on either side of the peak at which the z-scored signal crossed the mean. Interneuron spike density was not used for event detection but was analyzed relative to detected spike density events.

### Calculation of spike density event rate over time since stopping

Spike density event (SDE) rates were computed as a function of time following reward consumption onset. For all rewarded stops, SDEs were counted in non-overlapping 0.5 s time bins and converted to rates by dividing by the bin duration (0.5 s). Rates were calculated over the interval 0–10 s, where time 0 was defined as the moment the rat first came within 5 cm of the reward well while moving at <3 cm/s. All rewarded stops were included in the analysis regardless of duration. If the rat did not remain stationary for the full 10 s, time bins after the rat exited the reward zone or exceeded the speed threshold were set to NaN. For each trial, the resulting SDE rate time course was smoothed by convolution with a Gaussian kernel (0.5 s standard deviation). Figures report the mean ± SEM across trials. Sharp-wave ripple event rates were computed using the same procedure.

### Place fields

The rat’s position and speed at each spike time were computed through linear interpolation (*interp1* in MATLAB). Place fields were calculated from periods of movement (speed exceeding 5 cm/s). Positions were binned into 2 × 2 cm square bins (open field). Unsmoothed rate maps were computed as the number of spikes per bin normalized by occupancy. Place fields were defined as contiguous bins (minimum of 15 bins) with a firing rate greater than 20% of the peak firing rate, where the peak firing rate was > 1 Hz. Unvisited bins were set to zero, and the raw maps were convolved with a 2D isotropic Gaussian kernel (8 cm standard deviation). In Fig. 1, we examined the development of place fields over trial. For this analysis, place fields were constructed using the spiking and positional data up until each trial, rather than over the entire session.

### Bayesian decoding of position

The posterior probability (*P*) of the animal’s position (pos) is given by Bayes’ rule (assuming Poisson firing statistics, independence between neurons, and a uniform prior over position^30^). The posterior probability of the animal’s position (pos) across *L* total position bins given a time window (*τ*) containing neural spiking (spikes) is

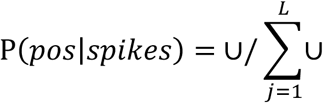

and

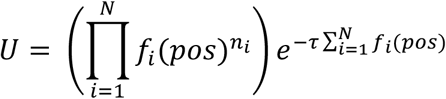

where *f*_*i*_(pos) is the position tuning curve of the *i*^*th*^ unit, and *n*_i_ is the number of spikes emitted by the *i*^*th*^ neuron in the time window (*τ*). The posterior probability was computed for all spatial bins *x*_*j*_, where 1≤j≤L and L is the total number of spatial bins.

During movement, position was estimated from non-overlapping 400 ms windows and compared to the true position to determine the decoding error. Sessions for which the average decoded positional error exceeded 3 cm were discarded from further analysis. During immobility, position was estimated in smaller temporal windows (20 ms overlapping by 5 ms). The posterior probability over position was computed during each spike density event. Temporal bins in which the peak posterior probability was <0.01 were removed.

Posterior center-of-mass (COM): Let *P*_*j*_ = *P*(*x*_*j*_|spikes), where x_j_ =(x_j_,y_j_) denotes the coordinates of the *j*^*th*^ spatial bin. The posterior *center-of-mass* (COM) at a given time was defined as

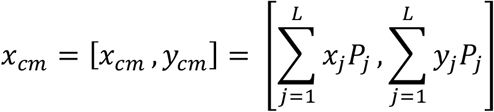

where *L* is the total number of spatial bins.

The distance covered by a replay event (“net displacement”) was defined as the Euclidean (L2) norm of the displacement between the first and last posterior COM estimates, 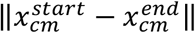. Instantaneous replay speed was defined as the Euclidean distance between consecutive COM estimates divided by the elapsed time between those estimates:

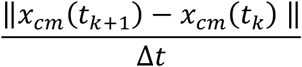

### Replay–past avoidance

Angular displacement from the rat’s past path was computed as described previously^8^. For each stopping period (reward consumption at either the Goal or Non-Goal well), we defined the rat’s past path as the 180 cm trajectory preceding reward consumption onset. For each replay event, a series of concentric circles (radii 2–180 cm, in 2 cm increments) was centered on the rat’s position at replay onset. For each circle, we computed the minor arc formed by its intersections with the replay trajectory and the rat’s preceding path. Angular displacements from circles with radii <12 cm (proximal) and >12 cm (distal) were averaged separately. Thus, each replay event yielded two values: the mean angular displacement relative to the past path at proximal distances and at distal distances. When multiple intersections occurred between a circle and either trajectory, the intersection closest in time to the replay was used.

### Replay–replay avoidance

To quantify replay–replay avoidance, we computed the angular displacement between pairs of replay events occurring within the same stopping period, as described previously^8^. Concentric circles (2–180 cm, in 2 cm increments) were centered on the rat’s position at the onset of the first replay event. For each circle, the minor arc formed by its intersections with the two replay trajectories was computed. Angular displacements were averaged separately for radii <12 cm (proximal) and >12 cm (distal), yielding two angular displacement measures per replay pair. Only locally initiated replays (those beginning within 12 cm of the rat’s position) were included. The mean angular displacement between replay pairs was plotted as a function of the time elapsed between them (Δt). Replay pairs were binned in 400 ms bins of Δt, advancing in 100 ms steps, and the mean angular displacement was computed for each bin. Statistical significance was assessed using a shuffle procedure in which Δt values were randomly permuted across replay pairs. For each of 5,000 iterations, Δt values were permuted and the Δt–angular displacement relationship was recomputed, generating a null distribution for each time bin. A bin was considered significant if the observed mean angular displacement fell outside the 2.5th–97.5th percentile range of the shuffled distribution.

### Replay displacement and speed as a function of the starting distance from the rat

Replay properties were quantified as a function of the distance between the rat and the start of each replay. For each replay event, we measured the start-to-end trajectory displacement (“net displacement”) and speed (“net speed”), and binned these measures according to the distance between the rat and the replay start location (2 cm bins from 0 to 24 cm). Within each bin, the mean and standard error of the mean (SEM) of replay displacement and speed were computed. Analyses were performed separately for replays occurring during Goal stops and Non-Goal stops. To ensure adequate sampling of distances, we pooled the data from all animals and sessions for this analysis (Fig. 2e-f).

### Quantification of remote replay properties

#### Speed

We asked whether nonlocal replay speed differed as trajectories approached a Goal well versus a Non-Goal well (Fig. 2h). We restricted analysis to replay events occurring within Non-Goal stops. For each decoding frame within a spike density event, we calculated the instantaneous replay speed and the decoded representation’s distance to the nearest well. Local representations (<12 cm from the rat) were excluded. We then binned replay speed according to the decoded representation’s distance from the nearest well (2 cm bins, 0 to 12 cm). To ensure adequate sampling of distances, we pooled the data from all animals and sessions for this analysis. Additionally, for each session we obtained the mean speed of remote replay representations within 12 cm of the Goal or a Non-Goal well (Fig. 2i).

#### Stopping probability

We asked whether replay stopping probability was higher at the Goal well compared to other locations in the following manner. Analyses were restricted to replay events occurring during Non-Goal stops. For each Non-Goal well A, we computed the conditional stopping probability at every other well T (the Goal well and remaining 7 Non-Goal wells) as: *P*(stop at T | entered T), defined as the number of replays that terminated within 12 cm of T divided by the number of replays that entered within 12 cm of T. To qualify as an entry, a replay trajectory had to start outside a 12 cm radius of the well and subsequently enter within 12 cm of the well. For each session, stopping probabilities were first computed separately for replay events originating from each Non-Goal well A, and then averaged across origin wells. We then compared, across sessions, stopping at the Goal well versus the mean stopping probability at Non-Goal wells.

#### Goal well overrepresentation

We asked whether the Goal well was overrepresented in remote replay during reward consumption at Non-Goal wells. For each session, we pooled all replay events occurring during Non-Goal stops and excluded local representations by removing decoded positions within 12 cm of the animal’s actual location. We then computed the mean nonlocal posterior probability within 12 cm of each of the 9 wells, yielding a measure of total nonlocal posterior probability per well for that session. Across sessions, we compared posterior density near the Goal well versus the average Non-Goal well.

### Identification of monosynaptically coupled pyramidal-interneuron pairs

Putative monosynaptic pyramidal–interneuron (Pyr–Int) connections were identified using spike-time cross correlogram (CCG) analysis computed over the entire behavioral session, including periods of locomotion. For each Pyr–Int pair, a CCG was computed using 0.4 ms bins. To account for slow rate co-modulation between neurons, a slow baseline expectation (*λ*_*slow*_) was estimated for each CCG by convolving the raw CCG with a Gaussian kernel (*σ* = 10 ms). The resulting smoothed CCG provided an estimate of the expected spike counts from slow rate co-modulation.

Pairs were classified as monosynaptic if a) the peak CCG count within the monosynaptic window (0.8-2.8 ms; positive lags, Pyr leading Int) exceeded the baseline expectation by more than 3 standard deviations (peak > *λ*_*slow*_ + 3√*λ*_*slow*_) and b) the total excess spike count above *λ*_*slow*_ within the monosynaptic window was at least twice that in the matched negative-lag window (-2.8 to -0.8 ms), ensuring directional asymmetry consistent with a Pyr–Int connection^15^.

Transmission probability (P_trans_) for each significant monosynaptic pair was quantified as:

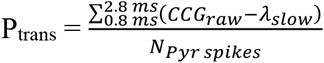

where the numerator represents the total excess spike count within the monosynaptic window (the summed excess spike count above λ_slow) and the denominator is the total number of pyramidal reference spikes during the session. This measure estimates the probability that a pyramidal spike is followed by an interneuron spike within the monosynaptic latency window.

For each putative monosynaptic pair, we determined whether the pyramidal neurons’s spatial firing rate map peaked near the Goal well or a Non-Goal well. The peak firing location was defined as the spatial bin with maximal firing rate, and neurons were classified as Goal-related or Non-Goal tuned if this peak fell within 12 cm of the Goal well or a Non-Goal well, respectively. For each session, we computed the mean transmission probability across significant Pyr–Int pairs in which the pyramidal neuron fired maximally near the Goal well and the mean transmission probability across significant pairs in which the pyramidal neuron fired maximally near any Non-Goal well. These transmission probabilities were then compared across sessions using a Wilcoxon signed-rank test.

### Simulations of replay sequences

Hippocampal sequences were simulated using a single-bump continuous attractor network (CAN) where the movement of the bump was driven by spike-frequency adaptation^8,10,35–37,39,45–47^. Given a summed input current to the *i*^*th*^ neuron, the instantaneous firing rate of the neuron was *f*(*I*_*i*_(*t*)), with the neural transfer function given by 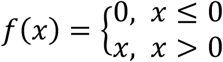.

Based on this time-varying input, neurons fired spikes according to a Poisson point process with a coefficient of variance of 1.

The input rate for the *i*^*th*^ neuron was a combination of internal and externally-derived inputs that were phase modulated in time:

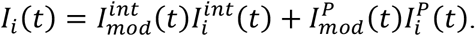

The “internal” input was given by

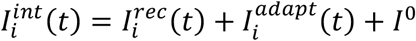

where 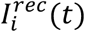 is the recurrent input derived from other neurons (see below), 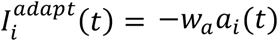 the adaptive inhibitory input (*W_a_* is the strength of depression) which models the effects of slow calcium-dependent potassium currents^39^, and *I*^*0*^ is a small positive constant bias common to all neurons. The recurrent input was: 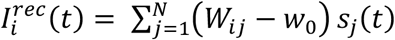,

where *W*_*ij*_ are the excitatory recurrent weights, *w*_0_ is the strength of recurrent inhibitory feedback, and *N* is the number of neurons. To specify the recurrent weights, neurons were organized into a 1D periodic array in the neural sheet, where the location of the *i*^*th*^ neuron was given by *x*_*i*_. Let *W*_*ij*_ be a set of translation-invariant symmetric weights with Gaussian shape that depend on the distance between neurons in the neural sheet:

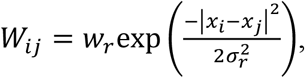

where *w*_*r*_ and *σ*_*r*_ control the strength and spatial extent of the connectivity, respectively. The synaptic activation dynamics for the *i*^*th*^ neuron, *s*_*i*_(*t*), was given by

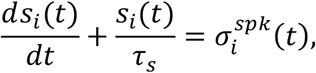

where *τ*_*s*_ is the synaptic time constant and

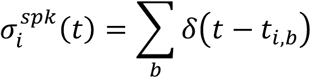

is the cell’s spike train (*t*_*i,b*_ specifies the time of the *b*^*th*^ spike of the neuron and the sum is over all spikes of the neuron). The adaptation dynamics for the *i*^*th*^ neuron, *a*_*i*_(*t*), was given by

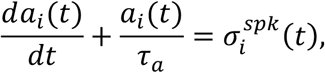

where *τ*_*a*_ is the time scale of adaptation.

To generate the corrective place inputs, a rat trajectory was simulated, starting at the origin, by generating a speed profile that was a sinusoid with period *T*_*transition*_ seconds but modified so that the rat dwelled in the rest state (zero velocity) for *T*_*rest*_ seconds and in the maximum velocity state for *T*_*run*_ seconds. This speed profile was then integrated to generate a spatial trajectory. The place input into the *i*^*th*^ neuron was given by a 1D Gaussian centered on the rat’s location at time *t* and width *σ*_*rat*_ and amplitude *w*_*rat*_:

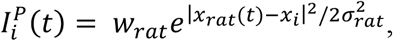

where *x*_*rat*_(*t*) is the rat’s position at time *t*

Lastly, to mimic the transient suppression of sequences during the transition from running to stopping that is seen in real data, we defined transitions to and from running as the timepoints when the rat’s speed rose above or dipped below 3 cm/s. Next, we convolved an inverted Gaussian with a 1.5 s standard deviation around each transition point, then renormalized the curve such that it ranged between [0,1]. This curve was then used to modulate the strength of the adaptation, *W_a_*, over time.

#### Phase modulation of internal and place inputs

Both the internal and place inputs were modulated by a rectified sinusoid with a frequency that was a function of rat speed, and selected to approximate the frequency of theta sequences during run, and replay sequences during rest:

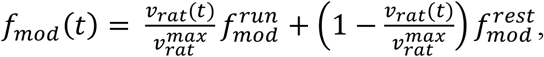

where 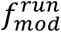 and 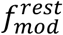 represent the maximum and minimum in the frequency of sequence generation and *v*_*rat*_(*t*) and 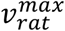 are the rat’s instantaneous and maximum running speeds, respectively (see below) (fig. S9). To introduce variability in sequence lengths, sequence periods were sampled iteratively from the distribution 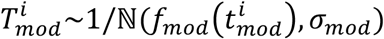, where ℕ is a normal distribution and 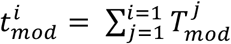 is the cumulative sum of all preceding sequence periods. The sequence modulation 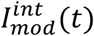 was then defined as follows: For the *i*^*th*^ sequence interval of length 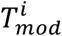 at time 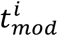,

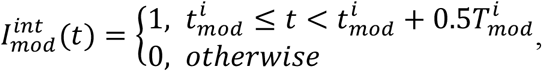

which amounts to a rectified sinusoid with a duty cycle of 50% of the sequence period. Likewise, the place inputs were also phase modulated:

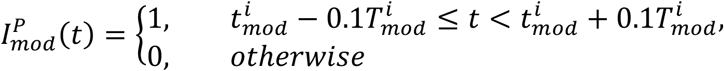

which amounts to a rectified sinusoid with a duty cycle of 20% of the sequence period, but also shifted in time in order to allow the place inputs to occur briefly around the time of the release of the internally-driven dynamics of the network.

#### Classification of sequences, replay–past avoidance, and replay–replay avoidance

Sequences were extracted from the dynamics of the model by tracking the center of mass of the activity bump during the “internal” phase of each sequence cycle. A sequence was required to activate at least 3 neurons, and was classified as extending either behind the animal (“reverse”, retracing the animal’s most recent path), or ahead of the animal (“forward”). Replay-past avoidance was defined as the difference between forward replay rate and reverse replay rate and was calculated over the stopping period. To compute replay-replay avoidance, we examined the frequency with which two sequences represented “Opposite” content as a function of the elapsed time between them (Δt). “Opposite” pairs consisted of one forward and one reverse replay. “Same” pairs consisted of either two forward or two reverse replays. We considered all pairs of locally initiated replays occurring within 4 s of one another in the same stopping period. Replay pairs were binned according to their Δt (400 ms bins advancing by 100 ms). For each time bin we computed the proportion of replay pairs classified as “Opposite”.

#### Simulation parameters

Integration was by the Euler method with step size equal to 0.5 msec. Other network parameters were:

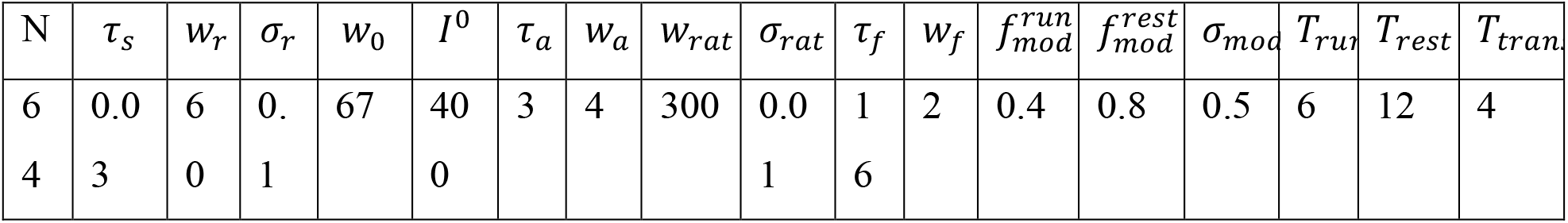

Temporal parameters are in units of seconds; amplitudes (*w*’s) are dimensionless; *I*^0^ is in units of spikes/sec.

### Statistical analysis

All statistical tests were two-tailed. In cases where data were not normally distributed, non-parametric statistical tests were used (Wilcoxon rank sum or Wilcoxon signed rank tests).

## Data Availability

All data will be made available upon request.

## Code Availability

All code will be uploaded to a repository upon publication.

## Acknowledgments

We thank all members of the Foster laboratory for discussions and feedback.

## Author Contributions

Conceptualization: C.S.M., D.J.F. Modeling: J.W. Methodology: C.S.M., J.W. Investigation: C.S.M., J.W. Visualization: C.S.M. Funding acquisition: C.S.M., D.J.F. Writing – original draft: C.S.M. Writing – review and editing: C.S.M., J.W., D.J.F.

