## Supplemental Figures for "Spatial goals trap hippocampal replay"

Supplementary Figures

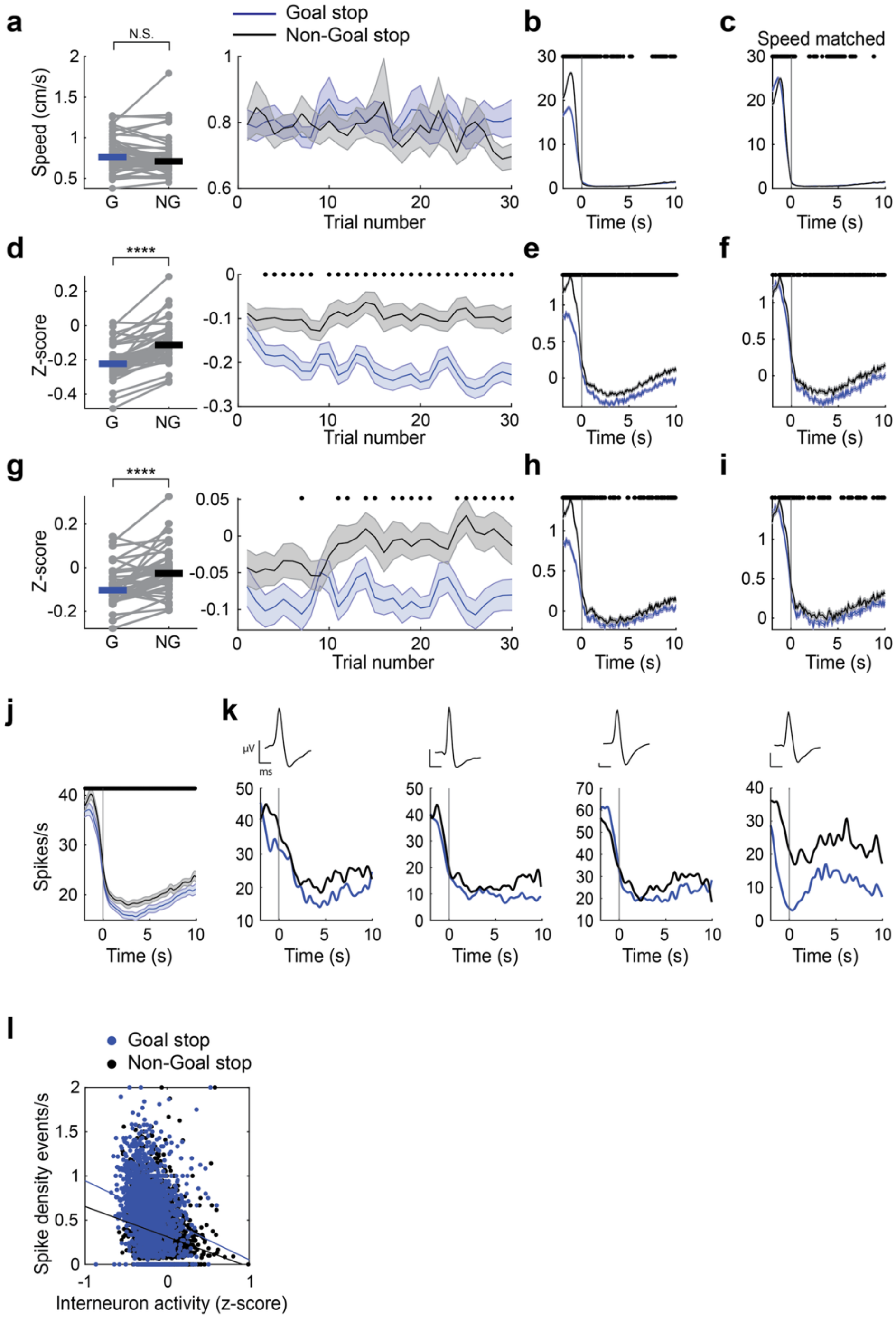


**Fig. S1. Interneuron firing rates during Goal and Non-Goal stops.**

**a.** Rat movement speed during Goal and Non-Goal stops. Left: average speed per session ($Z$=-1.8, $p$=0.071, signed-rank test). $N$=44 sessions. Right: movement speed during Goal and Non-Goal stops shown as a function of trial number (mean ± SEM).

**b.** Rat movement speed during the approach to and throughout the stop at Goal versus Non-Goal wells (N = 1960 Goal stops and 1939 Non-Goal stops). Dots indicate time bins for which $p$<0.05, rank-sum tests. Because rats tended to move more slowly during the approach to Goal versus Non-Goal wells, neural data were reanalyzed after downsampling stops to match running speeds during the 2 s approach period.

**c.** As in (b), but after downsampling stops to match running speeds during the 2 s approach period.

**d.** Z-scored interneuron population firing rates during Goal and Non-Goal stops, excluding periods within spike density events. Left: average interneuron population firing rates per session ($Z$=-5.6, $p$=2.4e-8). $N$=44 sessions. Right: interneuron population firing rates during Goal and Non-Goal stops shown as a function of trial number (mean ± SEM). Dots indicate bins for which $p<0.05$, rank-sum tests.

**e.** Interneuron firing rates during the approach to and throughout stopping at Goal versus Non-Goal wells ($N$=1960 Goal stops and 1939 Non-Goal stops). Dots show time bins for which $p<0.05$, rank-sum tests.

**f.** As in (e), but after downsampling trials to match running speeds during the 2 s approach period ($N$=1156 Goal and 1156 Non-Goal stops). Note that the reduction in inhibition at Goal stops is maintained after matching pre-stop speeds between conditions.

**g.** Z-scored interneuron population firing rates during Goal and Non-Goal stops, including periods within pyramidal spike density events. Left: average interneuron population firing rates per session ($Z$=-4.7, $p$=2.7e-6). $N$=44 sessions. Right: interneuron population firing rates during Goal and Non-Goal stops shown as a function of trial number (mean ± SEM). Dots show trials for which $p<0.05$, rank-sum tests. Note that the reduction in inhibition at the Goal is partially mitigated in this analysis because many interneurons participate in spike density events, which occur more frequently at the Goal.

**h.** Interneuron firing rates during the approach to and throughout stopping at Goal versus Non-Goal wells ($N$=1960 Goal stops and 1939 Non-Goal stops). Dots show time bins for which $p<0.05$, rank-sum tests.

**i.** As in (h), but after downsampling trials to match running speeds during the 2 s approach period ($N$=1156 Goal and 1156 Non-Goal stops).

**j.** Raw interneuron firing rates during the approach to and throughout Goal and Non-Goal stops, excluding periods within pyramidal spike density events. For each session, the mean firing rate of each interneuron across Goal or Non-Goal stops was computed, followed by averaging across interneurons to generate Goal and Non-Goal traces. Plots here depict the mean ± SEM across sessions. $N$=44 sessions. Dots indicate time bins for which $p<0.05$, signed-rank tests.

**k.** Firing rates of four representative interneurons during Goal and Non-Goal stops (excluding periods within pyramidal spike density events) in one session. Top: waveforms. Scale bars: 0.33 ms, 50 µV. Although many individual interneurons showed firing patterns consistent with the population average, some interneurons (e.g., last example) exhibited distinct response profiles.

**l.** There was a significant negative correlation between interneuron activity outside of spike density events and spike density event rate across stopping periods ($N$=3899 stopping periods). Three outlier stopping periods were omitted from the plot for visualization but included in statistical analyses. The association was significant for both Goal and Non-Goal stops (Goal: $r(1958)$=-0.23, $p$=7.6e-26; Non-Goal: $r(1937)$=-0.22, $p$=2.4e-23; Pearson correlations). Blue and black lines show the lines of best fit for Goal and Non-Goal stops.

****$p<0.0001$

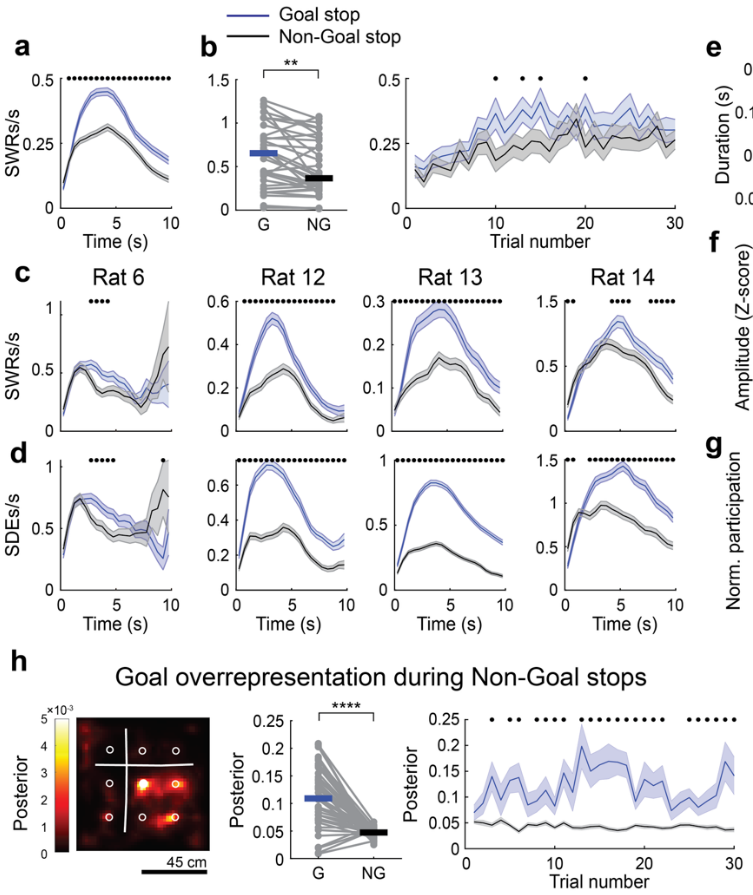


**Fig. S2. Reactivation event properties differ between Goal and Non-Goal stops**

**a.** As observed for spike density events (Fig 1), sharp-wave ripples (SWRs) occurred more frequently during Goal versus Non-Goal stops. Traces show SWR rate during reward consumption at Goal or Non-Goal wells ($N$=1503 Goal stops and 1486 Non-Goal stops). Dots indicate time bins for which $p$<0.05, rank-sum tests.

**b.** Left: Average SWR rate per session, computed separately for Goal and Non-Goal stops ($Z$=3.3, $p$=0.00010, signed-rank test). $N$=34 sessions with usable LFP data. Right: SWR rate during Goal and Non-Goal stops shown as a function of trial number (mean ± SEM). Dots indicate trials for which $p$<0.05, rank-sum tests.

**c.** Sharp-wave ripple event rate over time since stopping, shown separately for each animal. Rat 6: $N$=196 Goal stops and 193 Non-Goal stops; Rat 12: $N$=423 Goal stops and 417 Non-Goal stops; Rat 13: $N$=618 Goal stops and 612 Non-Goal stops; Rat 14: $N$=266 Goal stops and 264 Non-Goal stops.

**d.** Spike density event rate over time since stopping, shown separately for each animal. Rat 6: $N$=196 Goal stops and 193 Non-Goal stops; Rat 12: $N$=526 Goal stops and 518 Non-Goal stops; Rat 13: $N$=972 Goal stops and 964 Non-Goal stops; Rat 14: $N$=266 Goal and 264 Non-Goal stops.

**e.** The duration of spike density events was reduced during on Goal versus Non-Goal stops ($Z$=-4.1, $p$=4.2e-5, signed-rank test). $N$=44 sessions from 4 rats. Right plot shows duration as a function of trial number (mean ± SEM). Dots indicate trials for which $p$<0.05, rank-sum tests.

**f.** The amplitude of spike density events was greater on Goal versus Non-Goal stops ($Z$=4.7, $p$=3.2e-6, signed-rank test). $N$=44 sessions. Right plot shows amplitude as a function of trial number (mean ± SEM). Dots indicate trials for which $p$<0.05, rank-sum tests.

**g.** The percentage of neurons participating in an SDE normalized by SDE duration. Normalized participation was greater for SDEs during Goal stops versus Non-Goal stops ($Z$=2.9, $p$=0.0035, signed-rank test). Right plot shows normalized participation as a function of trial number (mean ± SEM). Dots indicate trials for which $p$<0.05, rank-sum tests.

**h.** The Goal location is overrepresented within SDEs during Non-Goal stops. *Left:* Heat map from one example session showing the average posterior probability across space during SDEs occurring at Non-Goal stops. Filled white circle: Goal well; open white circles: Non-Goal wells. Local representations (<12 cm from the animal) were excluded. *Middle:* Average posterior probability near the Goal (<12 cm) or near a given Non-Goal (<12 cm) during SDEs at Non-Goal stops, shown for each session (N = 44 sessions from 4 rats). Local representations (<12 cm from the animal) were excluded. *Right:* Remote posterior probability near the Goal or Non-Goal wells during SDEs at Non-Goal stops, plotted as a function of trial number (mean ± SEM). Dots indicate trials with $p < 0.05$, rank-sum tests.

**p<0.01, ****p<0.0001

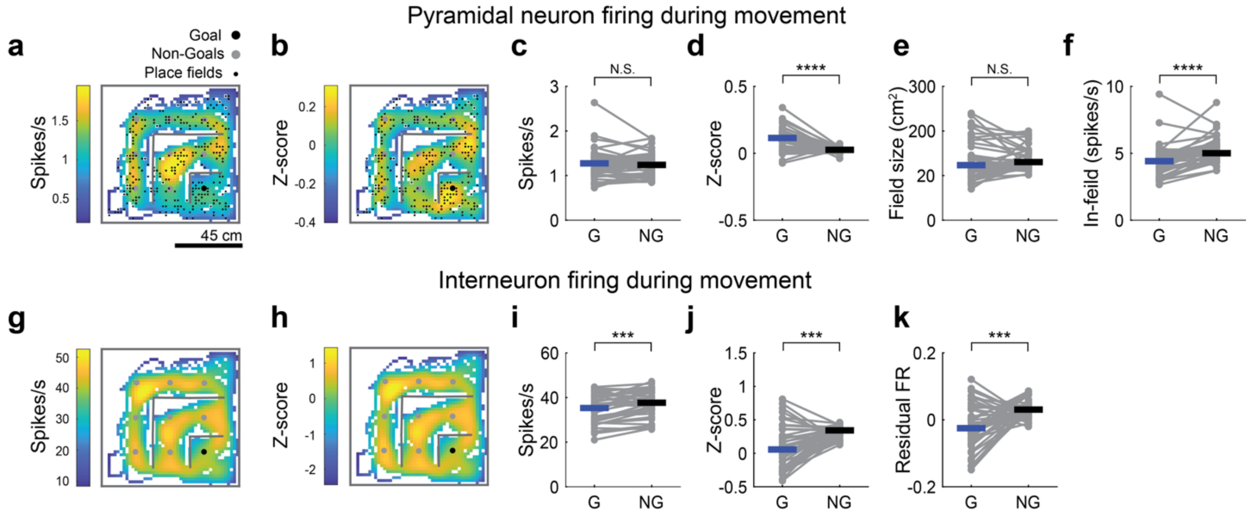


**Fig. S3. Place field overrepresentation and reduced inhibition near the Goal during movement**

- **a.** Example session showing the average pyramidal cell spatial firing rate map, obtained by averaging across cells. A small black dot indicates the center-of-mass of each detected place field. Spatial bins that were not traversed at >5 cm/s are shown in white. Consistent with previous reports using a similar task(Pfeiffer, 2022; Pfeiffer & Foster, 2013), the average firing rate around the Goal did not appear elevated relative to that around Non-Goal wells. However, place fields were more densely concentrated near the Goal (also see Fig. 1g).
- **b.** Example session showing the mean of all z-scored pyramidal cell spatial firing rate maps. Each cell's spatial firing rate map was z-scored across space prior to averaging. In this analysis, the overrepresentation of the Goal was substantially more apparent.
- **c.** Session-averaged mean firing rates near (within 12 cm of) the Goal or a Non-Goal, computed from the mean pyramidal cell spatial firing rate maps (as in a). Mean firing rates near the Goal did not significantly differ from those near Non-Goal wells ($Z$=1.5, $p$=0.14, signed-rank test). This apparent lack of overrepresentation is likely attributable to reduced running speeds near the Goal (Fig. S1), offsetting the increase in place field density (Fig. 1g). $N$=44 sessions from 4 rats.
- **d.** Pyramidal cells were more spatially tuned to the Goal. Plot shows session-averaged mean z-scored pyramidal cell firing within 12 cm of the Goal or Non-Goal wells. Z-scored firing rates near the Goal were higher than those near Non-Goal wells ($Z$=5.0, $p$=5.5e-7, signed-rank test). $N$=44 sessions. This analysis is consistent with the increased place field density at the Goal (Fig. 1J).
- **e.** Place field sizes did not differ between fields detected near a Goal and those detected near Non-Goal wells ($Z$=-1.1, $p$=0.26, signed-rank test). $N$=44 sessions. Fields were categorized as "Goal" if their center-of-mass was within 12 cm of the Goal, or "Non-Goal" if their center-of-mass was within 12 cm of a Non-Goal well.
- **f.** In-field firing rates were significantly reduced amongst place fields near the Goal relative to those near Non-Goal wells ($Z$=-4.3, $p$=1.8e-5, signed-rank test). $N$=44 sessions.
- **g.** Example session showing the mean of all interneuron spatial firing rate maps.
- **h.** Example session showing the mean of all z-scored interneuron spatial firing rate maps.
- **i.** Mean interneuron firing rates were reduced near (within 12 cm of) the Goal relative to near Non-Goal wells ($Z$=-3.6, $p$=0.00027, signed-rank test). $N$=44 sessions.
- **j.** Z-scored interneuron firing rates were also reduced near the Goal relative to near Non-Goal wells ($Z$=-3.5, $p$=0.00046, signed-rank test). $N$=44 sessions.
- **k.** Reduced interneuron firing near the Goal persisted after accounting for differences in running speed. For each interneuron, a linear model incorporating both position and running speed was fit to the data, and the contribution of running speed was removed to obtain residual firing attributable to position alone. Residual firing rates remained significantly lower near the Goal than near Non-Goal wells ($Z$=-3.4, $p$=0.00073, signed-rank test). $N$=44 sessions.

***$p$<0.001, ****$p$<0.0001

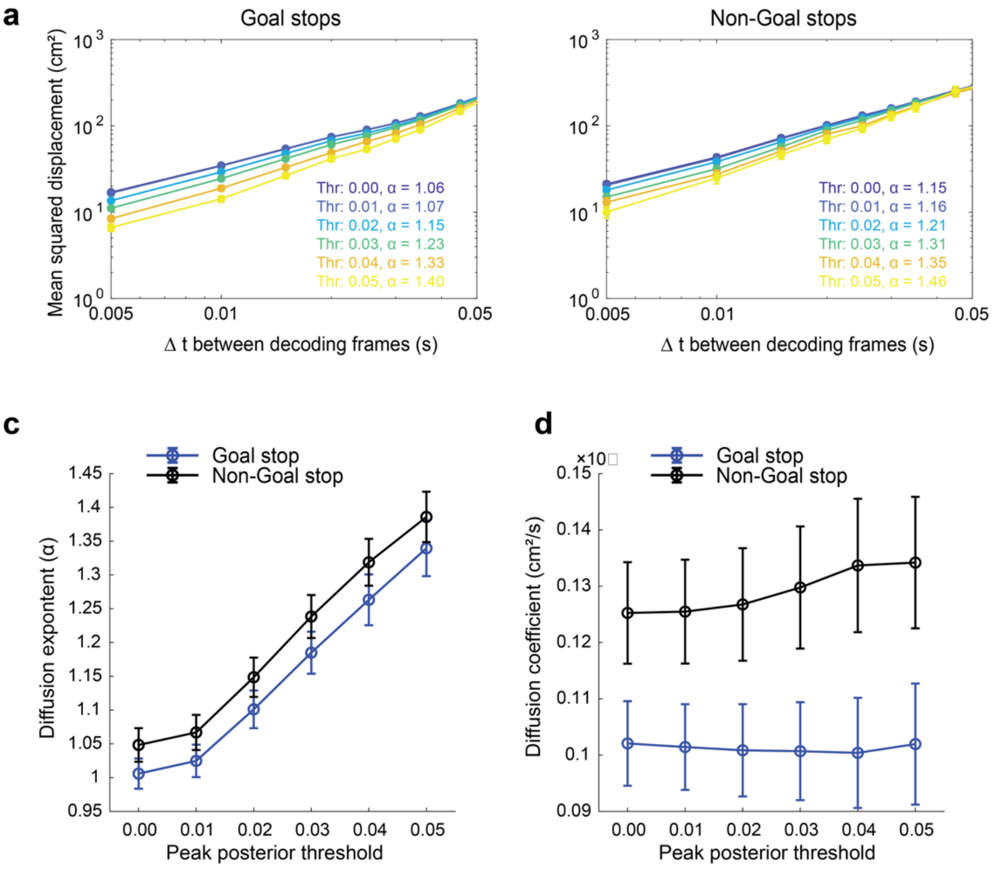


**Fig. S4. Diffusion dynamics of Goal- and Non-Goal-associated replays**

**a.** Example session showing the relationship between the time interval between decoding windows (Δt) and the mean squared displacement (MSD) of decoded positions during replay within Goal stops. Data are plotted on log-log axes. Replay diffusivity was estimated by fitting a power law ($MSD \propto \Delta t^{\alpha}$) using six replay inclusion criteria based on the peak posterior probability in each decoding bin. Thr0 indicates that all decoding frames were included, whereas Thr0.05 indicates that only decoding frames with a peak posterior probability >0.05 were included. The fitted diffusion exponent (α) is shown for each threshold.

**b.** As in (a), but for replay events occurring during Non-Goal stops in the example session.

**c.** The diffusion exponent (α) for replays at Goal or Non-Goal stops across peak posterior thresholds. Error bars show mean ± SEM exponents across 44 sessions. To determine whether replay diffusivity differed between Goal and Non-Goal stops across thresholds, we fit a linear mixed-effects model with Stop Type and Threshold as fixed effects and session as a random intercept. The diffusion exponent was significantly higher during Non-Goal versus Goal stops ($\beta$=0.048 ± 0.009 SE, $t(525)$=5.52, $p$=5.46e-8), indicating that replay trajectories were more superdiffusive during Non-Goal stops. The estimated exponent also increased with increasing posterior peak threshold, reflecting progressively stricter inclusion of decoded positions ($\beta$=0.071 ± 0.003 SE, $t(525)$=28.08, $p$=1.32e-106). In the main text and Fig. 2B we used a peak posterior threshold of 0.01.

**d.** The diffusion coefficient (D) for replays at Goal or Non-Goal stops across peak posterior probability thresholds. Error bars show the mean ± SEM coefficients across 44 sessions. A linear mixed-effects model with Stop Type and Threshold as fixed effects and session as a random intercept revealed that the diffusion coefficient was significantly higher during Non-Goal versus Goal stops ($\beta$=0.023 ± 0.0022 SE, $t(525)$=12.94, $p$=1.9e-33). This demonstrates that the spatial spread of replay trajectories is slower on Goal stops. In contrast to the diffusion exponent, the diffusion coefficient did not vary significantly with posterior probability threshold ($t(525)$=1.55, $p$=0.12).

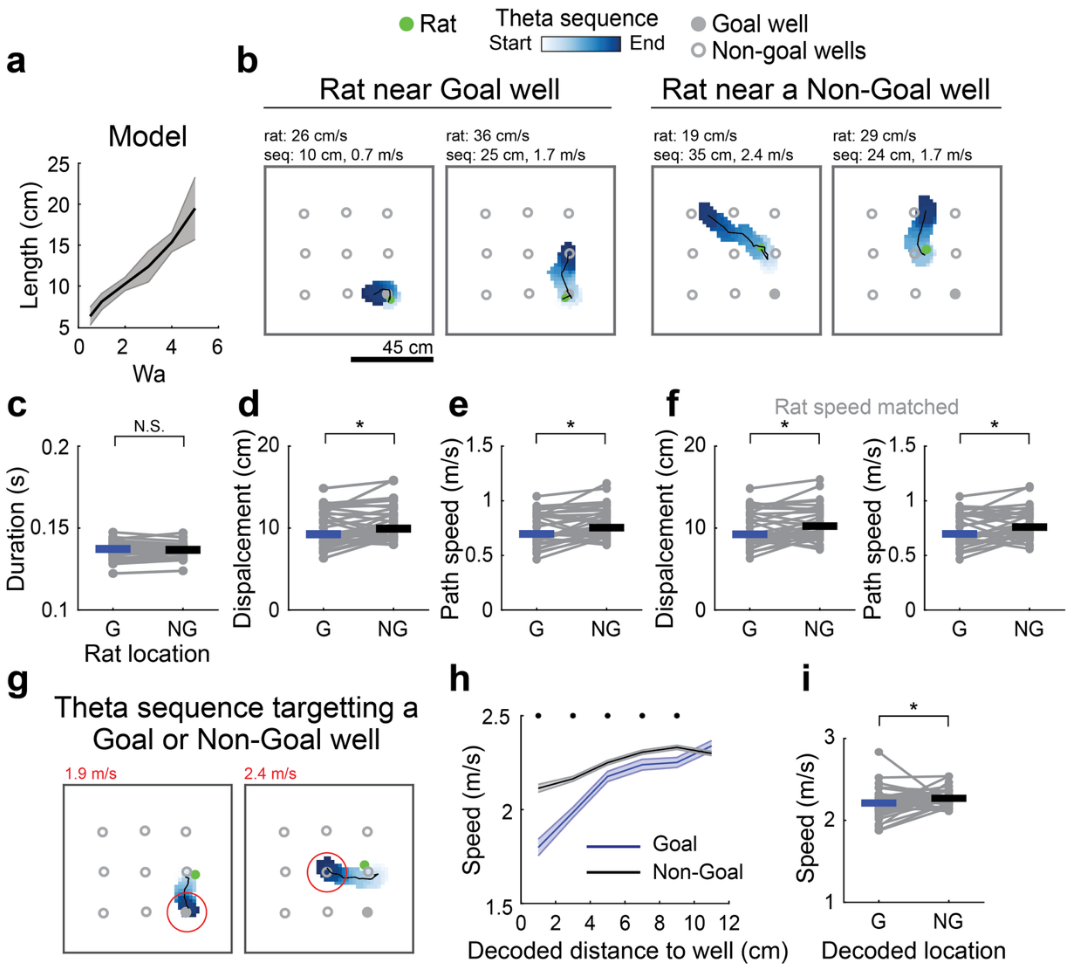


**Fig. S5. Slowed Goal representations within theta sequences**

**a.** In the network model, the length of sequences occurring during movement (i.e., theta sequences) increased with increasing adaptation strength ($r(4)$=0.998, $p$=6e-6, Pearson's correlation). Plot shows mean ± SEM replay length from 100 model runs each with $W_a$ = 0.5, 1, 2, 3, 4, or 5.

**b.** Example theta sequences detected when the rat was near (<12 cm from) a Goal or Non-Goal well. Rat speed, theta sequence length, and theta sequence speed are indicated above.

**c.** The duration of theta sequences occurring when the rat was near (<12 cm from) a Goal well did not significantly differ from those occurring when the rat was near a Non-Goal well ($Z$=-0.7, $p$=0.46, signed-rank test). $N$=34 sessions with simultaneous LFP recordings.

**d.** Theta sequences occurring when the rat was near a Goal well were shorter than those occurring when the rat was near a Non-Goal well (net displacement: $Z$=-2.5, $p$=0.012, signed-rank test). $N$=34 sessions.

**e.** Theta sequences occurring when the rat was near a Goal well were slower than those occurring when the rat was near a Non-Goal well (net speed: $Z$=-2.5, $p$=0.013, signed-rank test). $N$=34 sessions.

**f.** Differences in theta sequence length (left) and speed (right) persisted after downsampling sequences to match running speed distributions within each session, indicating that these effects are not explained by differences in the animal's movement speed (displacement: $Z$=-2.4, $p$=0.019; speed: $Z$=-2.5, $p$=0.023, signed-rank tests). $N$=34 sessions.

**g.** Two example theta sequences occurring when the animal was at a similar location, but terminating near either a Goal or Non-Goal well.

**h.** Instantaneous theta sequence path speed as a function of the decoded representation's distance from the nearest well. Traces show mean ± SEM instantaneous speeds pooled across 34 sessions and 4 rats. Blue: sequences approaching the Goal well; gray: sequences approaching a Non-Goal well. Local representations (<12 cm from the animal) were excluded. To control for differences in physical distance between the animal and decoded positions, sequences were restricted to those occurring when the animal was near an adjacent well (as in examples shown in f).

**i.** Instantaneous theta sequence speed was slower for representations near (<12 cm from) the Goal well relative to those near a Non-Goal well ($Z$=-2.4, $p$=0.016, signed-rank test). Local representations (<12 cm from the animal) were excluded from analysis. $N$=34 sessions.

*$p$<0.05, **$p$<0.01, ***$p$<0.001

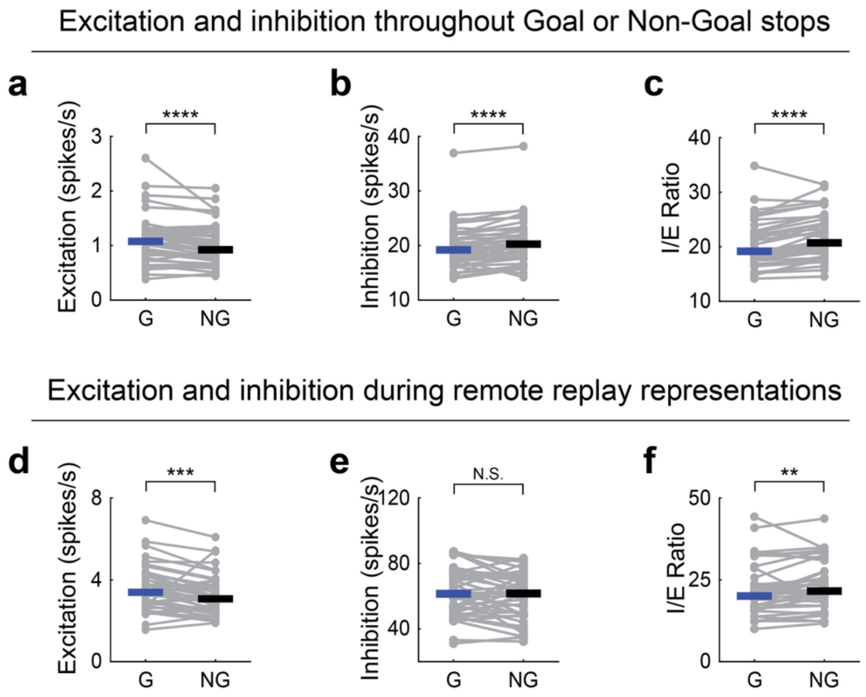


**Fig. S6. Reduced inhibition-to-excitation ratio at the Goal during behavior and remote replay**

- **a.** Excitation (mean firing rate of all pyramidal neurons) was higher during Goal versus Non-Goal stops ($Z$=4.2, $p$=2.4e-5, signed-rank test). This analysis included activity throughout the entire stopping period, including during spike density events. $N$=44 sessions.
- **b.** Inhibition (mean firing rate of all interneurons) was lower during Goal versus Non-Goal stops ($Z$=-4.2, $p$=3.1e-5, signed-rank test). This analysis included interneuron activity throughout the entire stopping period, including during spike density events. $N$=44 sessions.
- **c.** The inhibition-to-excitation ratio was lower during Goal versus Non-Goal stops ($Z$=-4.9, $p$=8.4e-7, signed-rank test). $N$=44 sessions.
- **d.** Excitation (mean firing rate of all pyramidal neurons) was higher during remote replay near (<12 cm) the Goal well than during remote replay near Non-Goal wells ($Z$=3.4, $p$=0.00066, signed-rank test). $N$=43 sessions containing at least one remote representation of each well type.
- **e.** Inhibition (mean firing rate of all interneurons) did not significantly differ between remote replay near (<12 cm) the Goal well and remote replay near Non-Goal wells ($Z$=1.7, $p$=0.096, signed-rank test). $N$=43 sessions containing at least one remote representation of each well type.
- **f.** The inhibition-to-excitation ratio was lower during remote replay representations near (<12 cm) the Goal well than during remote replay near Non-Goal wells ($Z$=-2.7, $p$=0.0061, signed-rank test). $N$=43 sessions containing at least one remote representation of each well type.

***$p$<0.001, ****$p$<0.0001
